# A comprehensive plasma metabolomics dataset for a cohort of mouse knockouts within the international mouse phenotyping consortium

**DOI:** 10.1101/624437

**Authors:** Dinesh K Barupal, Ying Zhang, Tong Shen, Sili Fan, Bryan S Roberts, Patrick Fitzgerald, Benjamin Wancewicz, Luis Valdiviez, Gert Wohlgemuth, Gregory Byram, YingYng Choy, Bennett Haffner, Megan R. Showalter, Arpana Vaniya, Clayton S Bloszies, Jacob S Folz, Tobias Kind, Oliver Fiehn

## Abstract

Mouse knockouts allow studying gene functions. Often, multiple phenotypes are impacted when a gene is inactivated. The International Mouse Phenotyping Consortium (IPMC) has generated thousands of mouse knockouts and catalogued their phenotype data. We have acquired metabolomics data from 220 plasma samples of 30 mouse gene knockouts and corresponding wildtype mice from IMPC. To acquire comprehensive metabolomics data, we have used liquid chromatography (LC) combined with mass spectrometry (MS) for detecting polar and lipophilic compounds in an untargeted approach. We have also used targeted methods to measure bile acids, steroids and oxylipins. In addition, we have used gas chromatography GC-TOFMS for measuring primary metabolites. The metabolomics dataset reports 832 unique structurally identified metabolites from 124 chemical classes as determined by ChemRICH software. The GCMS and LCMS raw data files, intermediate and finalized data matrices, R-Scripts, annotation databases and extracted ion chromatograms are provided in this data descriptor. The dataset can be used for subsequent studies to link genetic variants with molecular mechanisms and phenotypes.

**Data Set:** The dataset is available at the MetabolomicsWorkbench repository (accession ID: ST001154)

**Data Set License:** license under which the data set is made available (CC0).

## 1. Summary

Since 2000, the availability of the human genome sequence has fueled scientific investigations to link cellular functions and mammalian genetic variances[1]. Yet, for many genes, organismal functions are still unclear, hampering their use in clinical and translational approaches. Often, more than one function is affected by gene inactivation due to gene pleiotropy, which has also been observed in genome wide association studies (GWAS)[2–5]. Gene functions can be characterized on different biological levels from metabolite levels to cellular or whole-body level phenotype. GWAS catalogues such as the database of Genotypes and Phenotypes (dbGap) started associating various phenotypes with genetic variants[6], but such associations lack causal relationships. Here, animal models help chart molecular pathways from genetic variant to phenotype[7]. The International Mouse Phenotyping Consortium (IMPC) is a network of centers with expertise in mouse genetics and phenotyping. IMPC has established pipelines to generate knockout mice for over 7,000 genotypes and aim to cover all 20,000 protein coding genes in mice[8, 9]. The consortium has also generated mouse models for 360 diseases by creating 3,328 mouse knockouts[10]. The IMPC uses high throughput assays to measure phenotypes throughout the life of a knockout mouse and have successfully associated 974 genes with metabolic phenotypes and diseases[11]. Biomedical researchers can avail IMPC services to receive specific knockout biospecimens and search associated phenotype data using the mousephenotype.org website. All the IMPC generated data are publicly available at http://www.mousephenotype.org.

We here present metabolomic data attributed to mouse gene knockouts. Metabolic phenotypes are detailed by associating changes in metabolite levels (such as high cholesterol or low plasma uric acid) with genetic variants. Metabolism is dysregulated in many diseases. Up to 10% of all human genes are involved in regulating metabolism[12]. Several genes have well-characterized metabolic phenotypes. Currently, the IMPC measures only few metabolic endpoints such as body mass, plasma triglycerides, glucose tolerance, and basal blood glucose levels, warranting the need to expand their metabolic phenotype spectrum[11]. Over the past 20 years, metabolomics[13–15] has achieved an increased breadth and depth of analysis due to advances in sensitivity and accuracy of mass spectrometers with up to 900 identified metabolites measured in blood plasma[16].

In this data descriptor, we provide a comprehensive metabolomics dataset and a phenotype dataset for plasma specimens of 30 mouse knockouts and their normal wild type strains. Data were acquired by integrating three non-targeted assays (on primary metabolism, biogenic amines and complex lipids) with two targeted assays (oxylipins and a combined bile acids and steroids), using both GC-TOFMS and different LC-MS protocols.

## 2. Data Description

Raw GC-TOF MS and LC-MS mass spectra files are available at the NIH MetabolomicsWorkbench database (http://metabolomicsworkbench.org) (Data citation 1-7). Processed data matrices for all assays are provided in the data citation 8. The final metabolomics dataset is provided at (Data citation 9). Phenotype data for the mouse strains is provided at (Data citation 10). Data dictionary (Table 13), data matrix (Table S14), and sample metadata (Table S15) are provided in the supplementary section. Data file to sample label mapping is provided in the Table S10. The file also contains sample label to IMPC accession IDs so metabolite to phenotype data can be linked. Analysis sequences for each assay are provided in the Table S17 for checking if there were any batch effects or systematic error within the datasets. Annotation files are provided in the Table S3-S7. MRM transitions for the targeted assays are provided in the Table S8-S9. Processed results for each assay are provided in the Supplemental Table S11.

To ensure a high data quality, the following strategies were adopted while analyzing these samples: (a) use of internal standards mixture, (b) analysis of a quality control blood plasma samples, (c) analysis of blank samples to monitor carry-over and chemical artifacts including laboratory contaminants, (d) removal of multiple metabolites detections in different metabolomic platforms, (e) signal corrections using SERRF normalization for GC-TOF MS data, (f) removing compounds with > 50% missing values, (g) removing compounds with > 50% RSD technical variance, (h) using curated annotation databases to form a target list for peak intensity data processing, and (i) mapping peaks with compound identifiers and SMILES code for informatics analyses.

To show the technical reproducibility of the utilized LC-MS assays, RSD for peak heights of the internal standards were computed. Table 2 shows the RSDs values for these standards. Median RSDs for the detected compounds were 8% (GCMS), 11.5 % (CSH-POS), 13% (CSH-NEG), 12% (HILIC-POS) and 52% (HILIC-NEG). No batch effect was observed from the HILIC-POS, CSH-POS and CSH-NEG datasets. For HILIC-NEG, four batches were observed, and the signals were corrected using the median-batch normalization.

**Table 1.** Details of the mouse strains.

| **NCBI Gene ID** | **Gene Symbol** | **IMPC Line** | **Gene Description** |
| --- | --- | --- | --- |
| 235661 | Dync1li1 | K2P2 | Dynein Cytoplasmic 1 Light Intermediate Chain 1 |
| 71742 | Ulk3 | K2P2 | unc-51-like kinase 3 |
| 14380 | G6pd2 | KOMP2 | Glucose 6-phosphate dehydrogenase 2 |
| 29875 | Iqgap1 | KOMP2 | IQ motif containing GTPase activating protein 1 |
| 23980 | Pebp1 | KOMP2 | phosphatidylethanolamine binding protein 1 |
| 30939 | Pttg1 | KOMP2 | pituitary tumor-transforming gene 1 |
| 11947 | Atp5b | NorCOMM2 | ATP synthase, H<sup>+</sup> transporting mitochondrial F1 complex, beta subunit |
| 11972 | Atp6v0d1 | NorCOMM2 | ATPase H<sup>+</sup> Transporting lysosomal V0 Subunit D1 |
| 12567 | Cdk4 | NorCOMM2 | Cyclin Dependent Kinase 4 |
| 13361 | Dhfr | NorCOMM2 | Dihydrofolate reductase |
| 68421 | Lmbrd1 | NorCOMM2 | LMBR1 domain containing 1 |
| 18005 | Nek2 | NorCOMM2 | NIMA (never in mitosis gene a)-related expressed kinase 2 |
| 67963 | Npc2 | NorCOMM2 | NPC intracellular cholesterol transporter 2 |
| 19193 | Pipox | NorCOMM2 | Pipecolic acid oxidase |
| 19877 | Rock1 | NorCOMM2 | Rho-associated coiled-coil containing protein kinase 1 |
| 269378 | Ahcy | NorCOMM2 | S-adenosylhomocysteine hydrolase |
| 232345 | A2m | NorCOMM2 | alpha-2-macroglobulin |
| 230558 | C8a | NorCOMM2 | complement component 8, alpha polypeptide |
| 14420 | Galc | NorCOMM2 | galactosylceramidase |
| 26384 | Gnpda1 | NorCOMM2 | glucosamine-6-phosphate deaminase 1 |
| 15926 | Idh1 | NorCOMM2 | isocitrate dehydrogenase |
| 67096 | Mmachc | NorCOMM2 | methylmalonic aciduria cblC type, with homocystinuria |
| 17855 | Mvk | NorCOMM2 | mevalonate kinase |
| 76293 | Mfap4 | NorCOMM2 | microfibrillar-associated protein 4 |
| 54128 | Pmm2 | NorCOMM2 | phosphomannomutase 2 |
| 16922 | Phyh | NorCOMM2 | phytanoyl- CoA hydroxylase |
| 18817 | Plk1 | NorCOMM2 | polo-like kinase 1, serine/threonine protein kinase |
| 19248 | Ptpn12 | NorCOMM2 | protein tyrosine phosphatase, non-receptor type 12 |
| 24068 | Sra1 | NorCOMM2 | steroid receptor RNA activator 1 |
| 22631 | Ywhaz | NorCOMM2 | tyrosine 3-monooxygenase/tryptophan 5-monooxygenase activation protein |

**Table 2.**
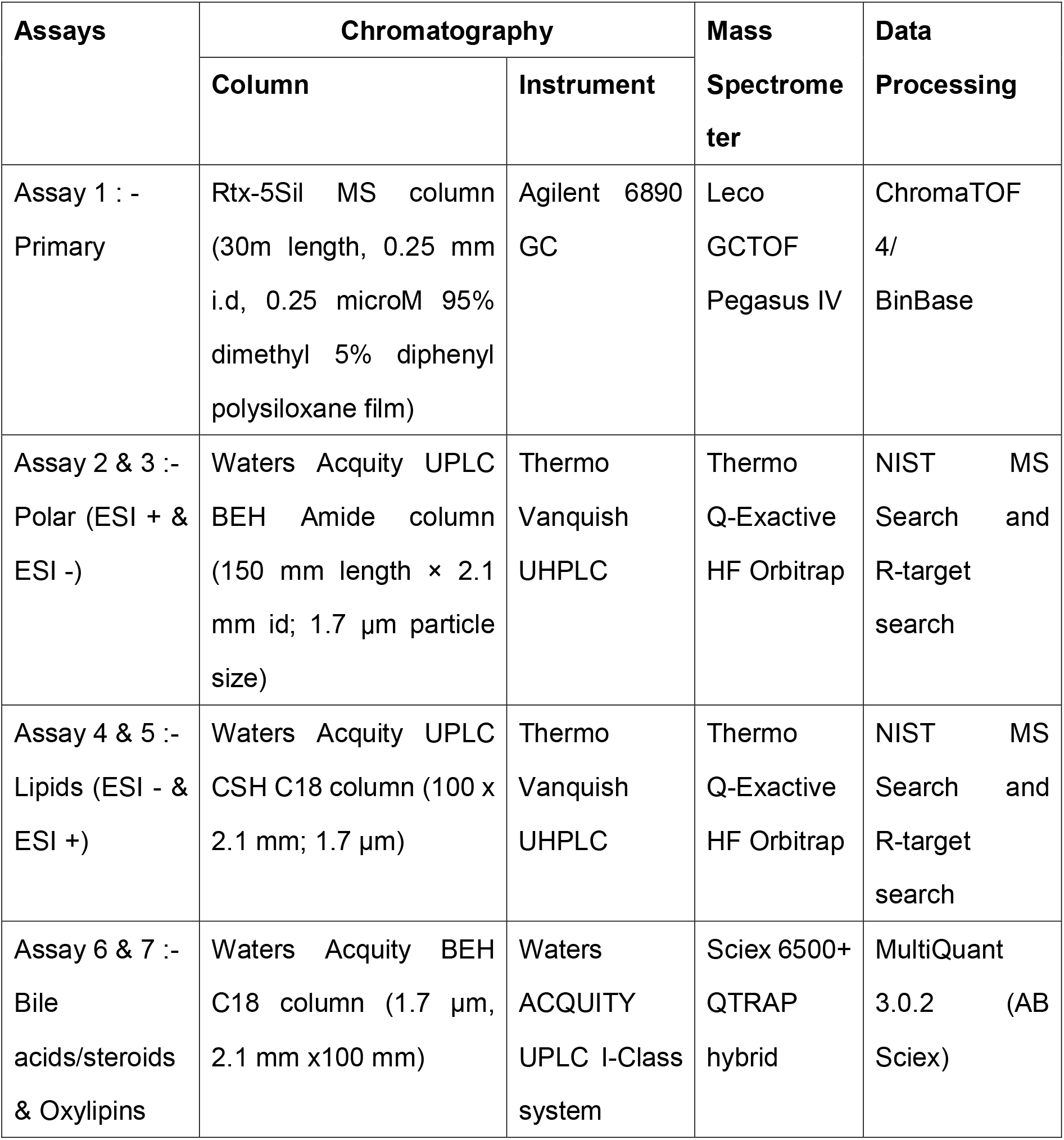
Overview of the Analytical Assays

| Assays | Chromatography |  | Mass | Data |
| --- | --- | --- | --- | --- |
|  | Column | Instrument | Spectrometer | Processing |
| Assay 1 : -<br>Primary | Rtx-5Sil MS column<br>(30m length, 0.25 mm<br>i.d, 0.25 microM 95%<br>dimethyl 5% diphenyl<br>polysiloxane film) | Agilent 6890<br>GC | Leco<br>GCTOF<br>Pegasus IV | ChromaTOF<br>4/<br>BinBase |
| Assay 2 & 3 :-<br>Polar (ESI + &<br>ESI -) | Waters Acquity UPLC<br>BEH Amide column<br>(150 mm length × 2.1<br>mm id; 1.7 µm particle<br>size) | Thermo<br>Vanquish<br>UHPLC | Thermo<br>Q-Exactive<br>HF Orbitrap | NIST MS<br>Search and<br>R-target<br>search |
| Assay 4 & 5 :-<br>Lipids (ESI - &<br>ESI +) | Waters Acquity UPLC<br>CSH C18 column (100 x<br>2.1 mm; 1.7 µm) | Thermo<br>Vanquish<br>UHPLC | Thermo<br>Q-Exactive<br>HF Orbitrap | NIST MS<br>Search and<br>R-target<br>search |
| Assay 6 & 7 :-<br>Bile<br>acids/steroids<br>& Oxylipins | Waters Acquity BEH<br>C18 column (1.7 µm,<br>2.1 mm x100 mm) | Waters<br>ACQUITY<br>UPLC I-Class<br>system | Sciex 6500+<br>QTRAP<br>hybrid | MultiQuant<br>3.0.2 (AB<br>Sciex) |

Targeted assays utilized a ten-point calibration curve to calculate molar concentrations of the target analytes. Values that did not pass the limit of quantification were not included in the data matrix, leading to many missing values. Table 3. Relative standard deviation of labelled internal standards for the LC/MS assays.

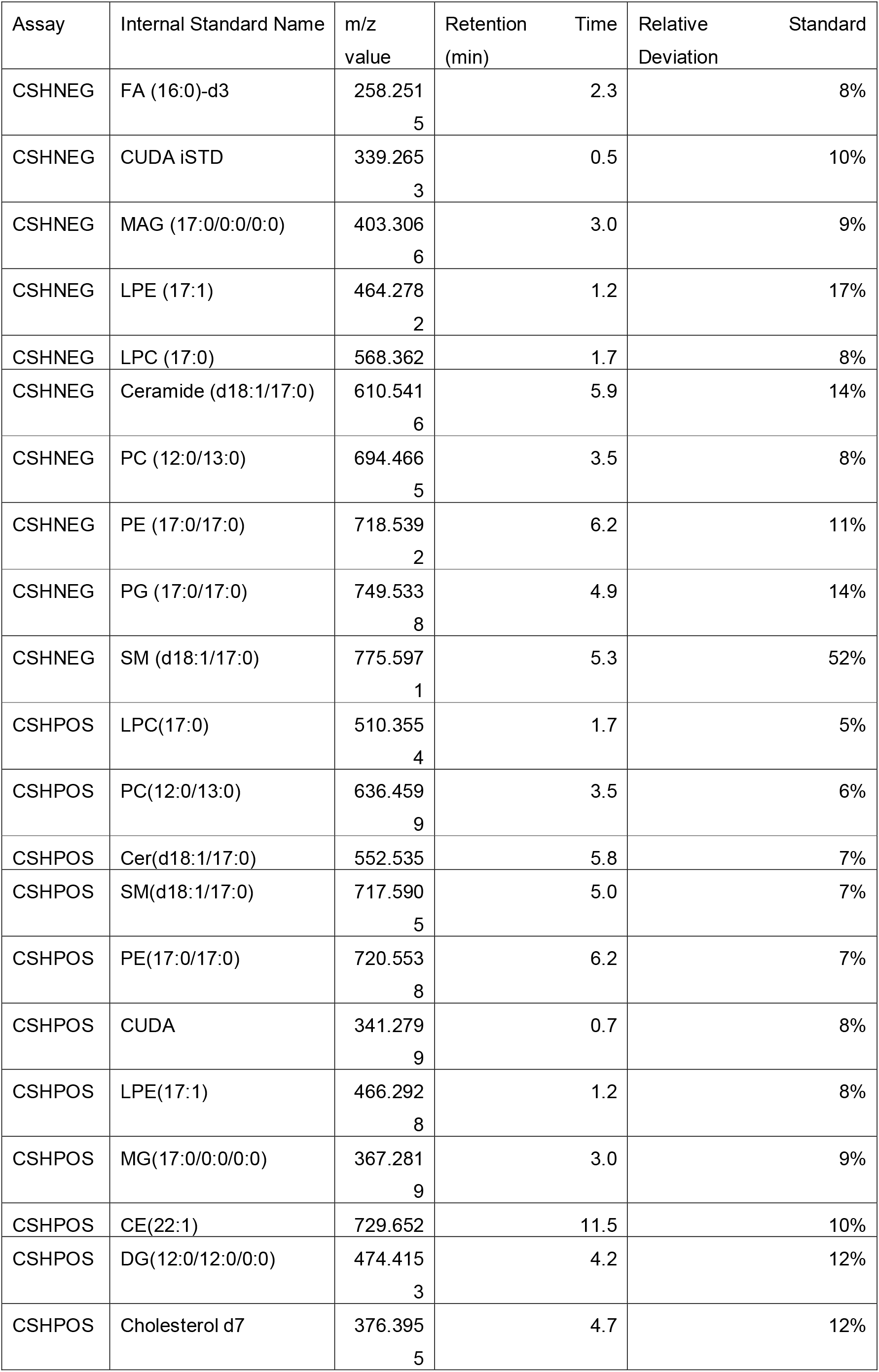
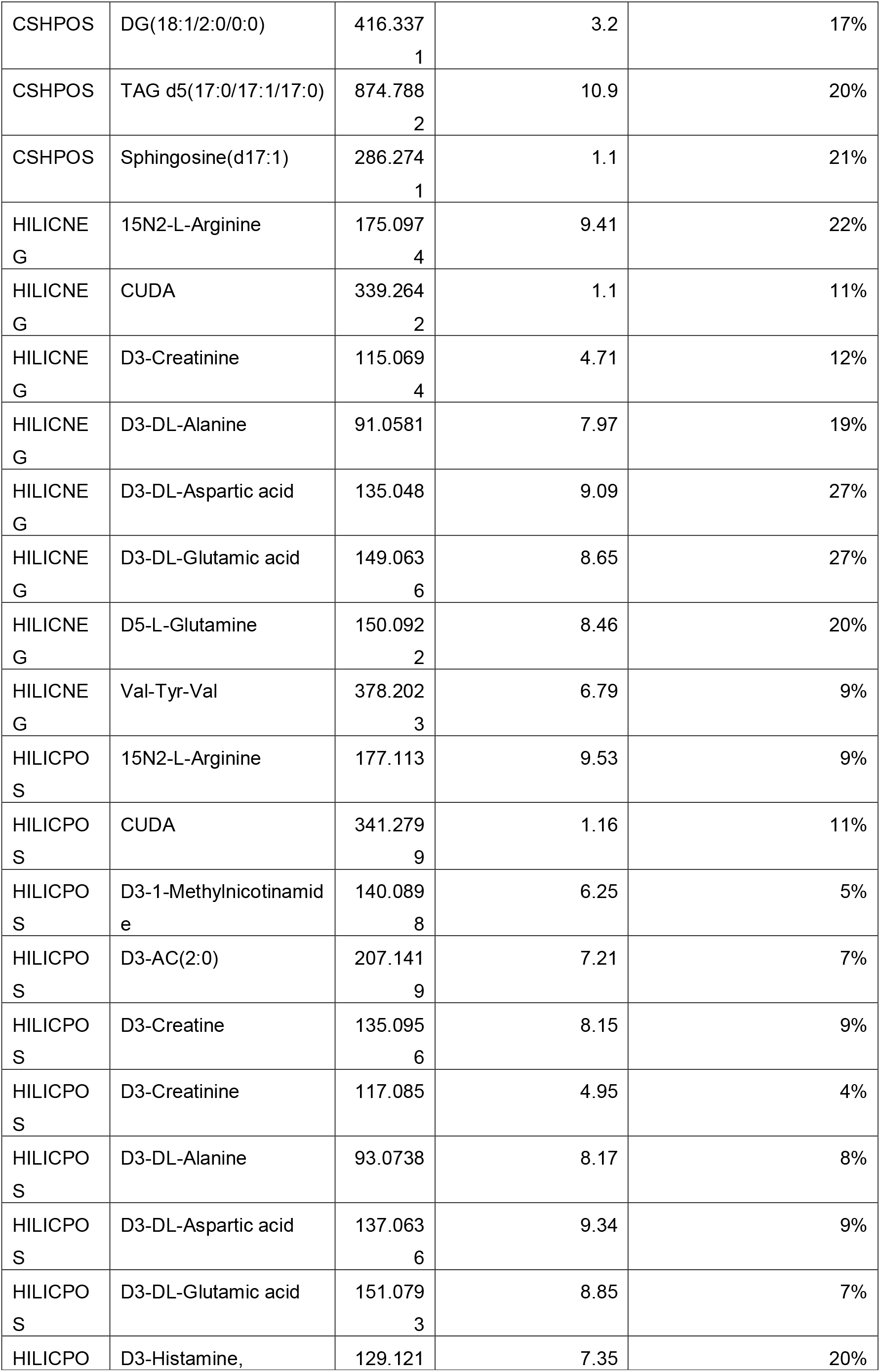
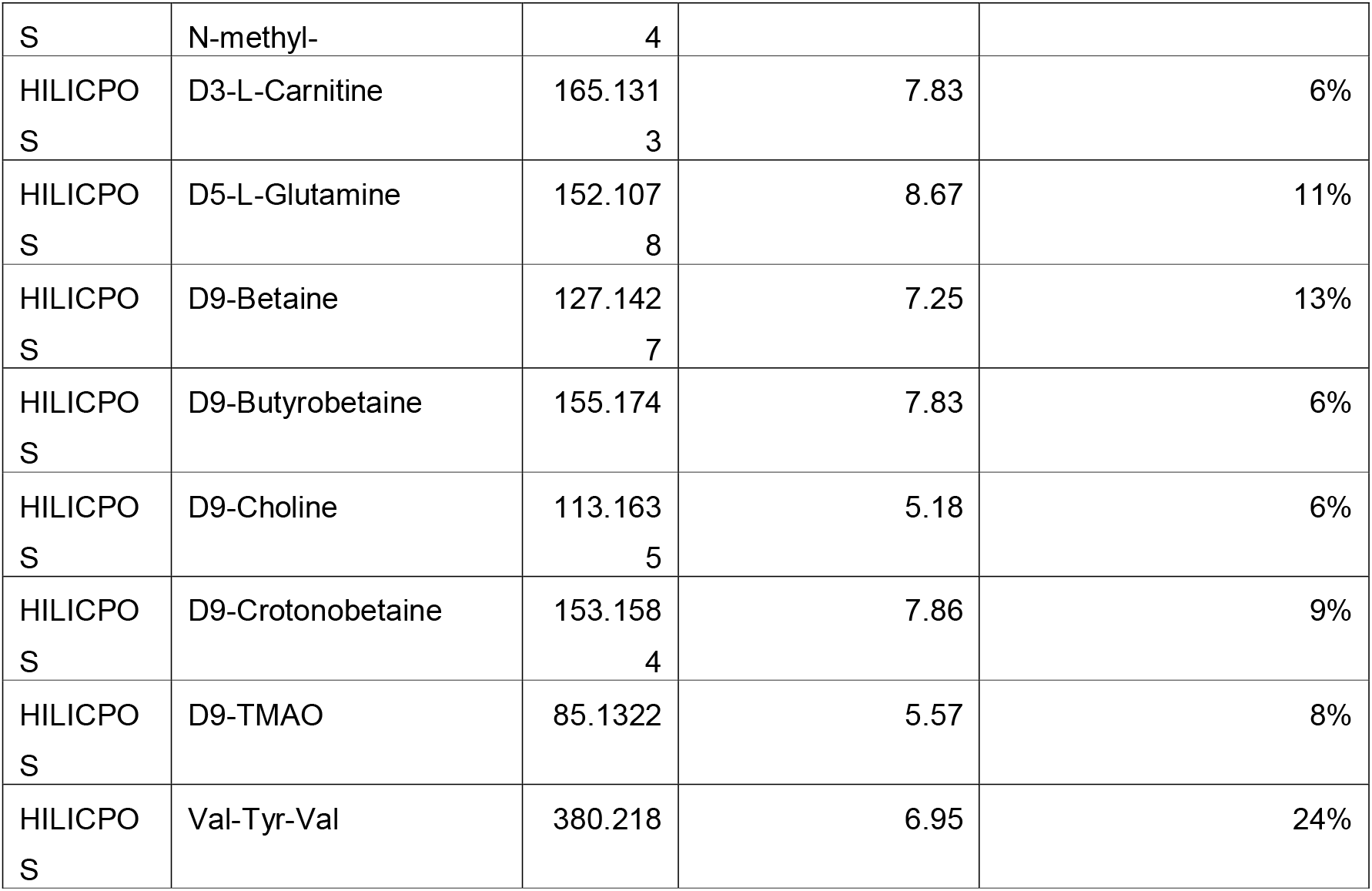

## 3. Methods

### IMPC consortium, mouse knockout selection and plasma samples

The International Mouse Phenotyping Consortium (www.mousephenotype.org) provided blood plasma samples for 30 knockouts (Table 1). For each knockout, three male and three female mice were selected, and a total of 40 wild-type mice were used to match the knockout strains. Plasma samples were shipped to the West Coast Metabolomics Center (WCMC; http://metabolomics.ucdavis.edu) on dry ice. Samples were stored at −80°C until analysis. Each sample was assigned a unique identifier according to the sampling date and time at three IMPC centers (See Table S15). Twenty additional human pool plasma samples (BioIVT, previously known as BioreclamationIVT) and up to 10 method blanks were analyzed with the mouse plasma samples for each analytical assay. Knockout genes were selected by sample availability, supposed effects on metabolism, and availability of a target assay in the proteomics core of the IMPC.

### Metabolomics facility

Metabolomics data for the mouse plasma were acquired using seven analytical assays using GC-MS and LC-MS platforms (Table 2). All LC-MS methods were performed using electrospray ionization (ESI). These assays are routinely used to generate metabolomics data at the WCMC for almost 30,000 samples per year, including many blood samples [13, 15, 17, 18]. The WCMC use large and validated lists of metabolite targets (Table S3-S9), large mass spectral libraries from the MassBank of north America (MONA available at http://massbank.us) to annotate novel compounds, standardized samples preparation and data acquisition methods, robust data processing using freely available MS-DIAL[19] and SERFF software[20], the BinBase mass spectral database[21] for covering over 150,000 GC-TOFMS samples analyzed over the past 15 years, and a variety of data analysis and interpretation tools, including statistics[22], pathway and network mapping[23] and metabolite enrichment analysis[24]. Figure 1 shows the overview of the metabolomics data generation and quality control workflow.

**Figure 1.**
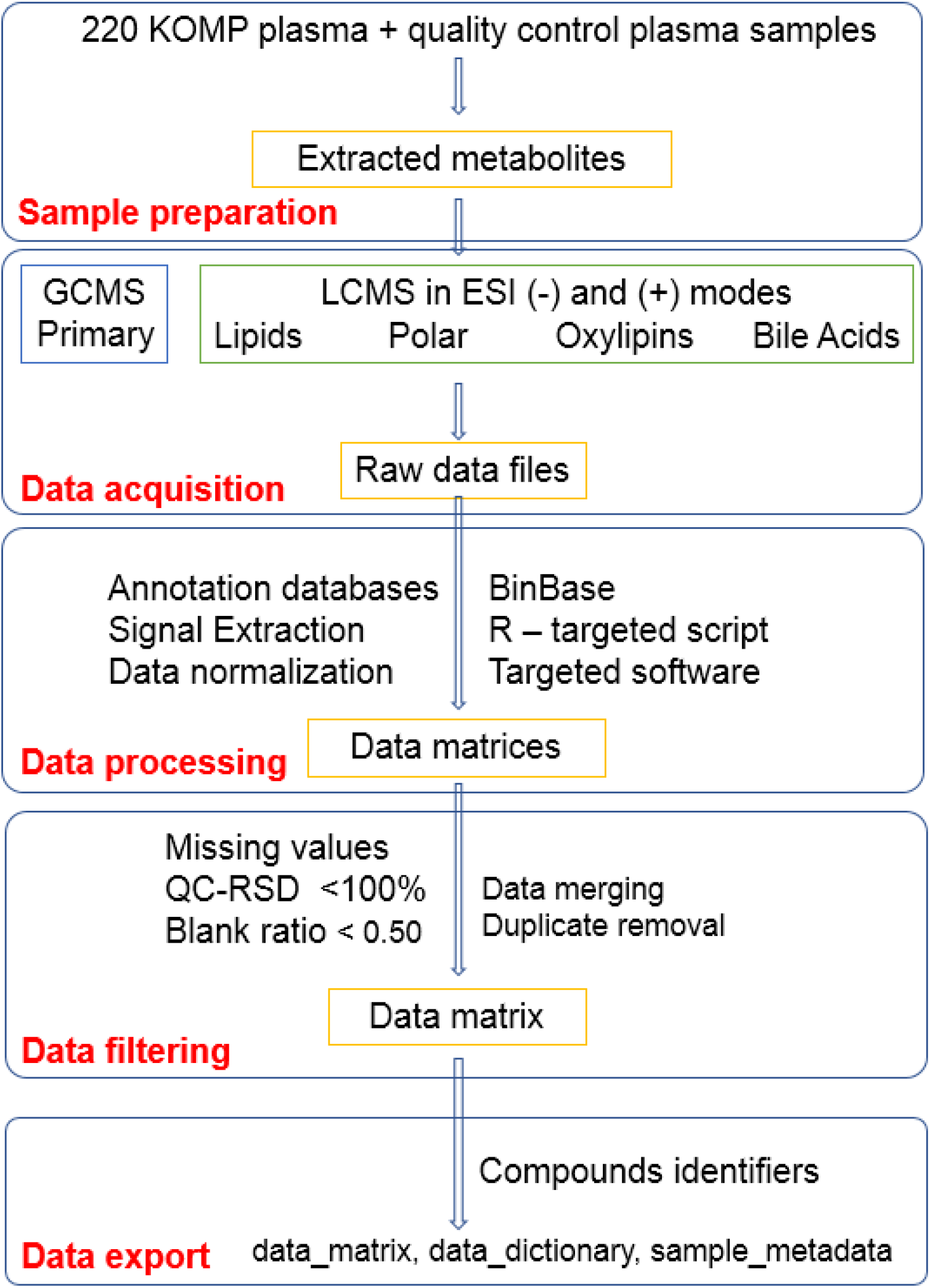
Overview of the metabolomics data generation and quality control workflow for 220 knockout mouse plasma (KOMP) samples. A less stringent relative standard deviation (RSD) and sample to blank ratio were used because the effect size of two or more show a major effect. As raw spectra files are provided for this study, a user can re-generate the data matrix with different thresholds. Abbreviation: GCMS-gas chromatography and mass spectrometry, LCMS – liquid chromatography and mass spectrometry and ESI – electrospray ionization.

### Annotation databases for untargeted metabolomics

#### Gas chromatography and mass spectrometry

Every acquired GC-TOFMS spectrum for blood specimens has been stored in the BinBase database for past 15 years at the WCMC. The database contain over 150,000 samples which can be queried through the BinVestigate web GUI (https://binvestigate.fiehnlab.ucdavis.edu/#/) for identified or unknown metabolites that are confidently detected in over 100 tissues and species[21]. The BinBase algorithm[25] utilizes this annotation database to generate a raw result data matrix (Table S11). The current BinBase annotation database is provided in supplementary Table S3 with 1205 annotated spectra for 588 unique compounds detected in biological samples.

#### Hydrophilic interaction liquid chromatography (HILIC) mass spectrometry

A database of target metabolites detected in HILIC-ESI-MS using both positive or negative electrospray mode are provided in supplementary Tables S4 and S5. This target database was generated by searching MS/MS spectra for blood specimens acquired in past three years against the NIST17 MS/MS, the LipidBLAST[26] and MoNA databases, in addition to a specific HILIC-retention time MS/MS mass spectral library of 1200 authentic standards[27]. For negative ESI mode, the HILIC-NEG annotation database yielded 107 identified compounds in the mouse plasma data set presented here using mass-to-charge (m/z), retention time (RT) and fragmentation spectra (MS/MS) match. An additional 45 compounds were verified by m/z and MS/MS fragmentation matches and one compound was confirmed by MZ and RT match. The abundance of this one compound was too low to trigger an experimental MS/MS event in data dependent MS/MS data acquisition methods. For the positive ESI mode, the HILIC-POS annotation database of the mouse plasma dataset presented here yielded 84 identified compounds that were confirmed by *Mm/z*, RT, and MS/MS matching, 86 compounds verified by *m/z*, and MS/MS data only, and 28 compounds were manually confirmed by *Mm/z* and RT match.

#### Charged surface hybrid liquid chromatography (CSH) and mass spectrometry

The CSH database for target mouse plasma lipids for positive and negative electrospray modes is provided in the supplementary table S6 and S7. The database is generated by searching MS/MS spectra for blood specimens acquired in past seven years against NIST17 MS/MS database and LipidBLAST mass spectral libraries. The CSH-NEG annotation database contains 215 verified lipids with MZ and MS/MS match; the CSH-POS annotation database contains 304 compounds with validated MZ and MS/MS match.

### Assay 1. Gas chromatography and mass spectrometry

#### Sample preparation

One milliliter of degassed, −20°C cold solvent mixture of acetonitrile (ACN):isopropanol (IPA):water (H2O) (3:3:2, v/v/v) was added to each 20 **μ**L mouse plasma aliquot. Samples were vortexed for 10 seconds, shaken for 5 minutes and then centrifuged for 2 minutes at 14,000 rcf (relative centrifugal force). Two 450 **μ**L supernatant aliquots was transferred to new tubes. To remove any excess protein, the supernatant was extracted with 500 **μ**L 1:1 acetonitrile:water and vortexed for 10 seconds, spun down for 2 minutes at 14,000 rcf. The supernatant was transferred to a clean tube and then dried down in a CentriVap concentrator. For derivatization, 10 **μ**L of methoxyamine hydrochloride in pyridine (40 mg/mL) was added to each sample and then shaken at 30°C for 90 minutes. Then 90 **μ**L of *N*-methyl-*N*-(trimethylsilyl) trifluoroacetamide (MSTFA, Sigma-Aldrich) was added for trimethylsilylation. C8-C30 fatty acid methyl esters (FAMEs) were added as internal standard (See Supplementary Table S18) for retention time correction. Samples were shaken for 30 minutes at 37°C. These derivatized samples were analyzed by GC-MS using a Leco Pegasus IV time of flight mass spectrometer. For more details see [28]

#### Data acquisition

An Agilent 6890 gas chromatography instrument equipped with a Gerstel automatic linear exchange systems (ALEX) which included a multipurpose sample dual rail and a Gerstel cold injection system (CIS). The CIS temperature program was: 50°C to 275°C final temperature at a rate of 12 °C/s and held for 3 minutes. Injection volume was 0.5 μL with 10 μL/s injection speed. Injection mode was splitless with a purge time of 25 seconds. Injector liner was changed after every 10 samples. Injection syringe was washed with 10 **μ**L of ethyl acetate before and after each run. A Rtx-5Sil MS column (30m length, 0.25 mm i.d, 0.25 microM 95% dimethyl 5% diphenyl polysiloxane film). An additional 10 m integrated guard column was used. Mobile phase was 99.9999% pure Helium gas with a flow rate of 1mL/min. GC temperature program was: held at 50°C for 1 min, ramped at 20°C/min to 330°C and then held for 5 minutes. A Leco Pegasus IV time of flight mass spectrometer was used to acquire data. The transfer line temperature between gas chromatograph and mass spectrometer was set to 280°C. Electron ionization at −70 V was employed with an ion-source temperature of 250°C. Acquisition rate was 17 spectra/second with a scan mass range of 85-500 Dalton (Da).

#### Data processing

Raw GC-TOF MS data files were preprocessed directly after data acquisition and stored as ChromaTOF-specific peg files, as generic txt result files and additionally as generic ANDI MS cdf files. ChromaTOF vs. 4.0 was used for data preprocessing without smoothing, 3 sec peak width, baseline subtraction just above the noise level, and automatic mass spectral deconvolution and peak detection at signal/noise (s/n) levels of 5:1 throughout the chromatogram. Results in.txt format were exported to a data server with absolute spectra intensities and further processed by a filtering algorithm implemented in the metabolomics BinBase database. The BinBase algorithm (rtx5) used the following settings: validity of chromatogram (10^7^ counts/s), unbiased retention index marker detection (MS similarity>800, validity of intensity range for high *m/z* marker ions), retention index calculation by 5^th^ order polynomial regression. Spectra were cut to 5% base peak abundance and matched to database entries from most to least abundant spectra using the following matching filters: retention index window ±2,000 units (equivalent to about ±2 sec retention time), validation of unique ions and apex masses (unique ion must be included in apexing masses and present at >3% of base peak abundance), mass spectrum similarity must fit criteria dependent on peak purity and signal/noise ratios and a final isomer filter. Failed spectra were automatically entered as new database entries if signal/noise ratios were larger than 25 and mass spectral purity better than 80%. All thresholds reflect settings for ChromaTOF v. 4.0. Quantification was reported as peak height using the unique ion as default, unless a different quantification ion was manually set in the BinBase administration software BinView. A quantification report table was produced for all database entries that were positively detected in more than 10% of the samples of this mouse knockout study. A subsequent post-processing module was employed to automatically replace missing values from the.cdf files. Prior to statistical analyses, data were filtered by combining multiple signals associated with each unique metabolite due to derivatization reactions. All metabolic signals were discarded if s/n>3 in comparison to blanks, or if replaced values were >3 the intensity of truly detected values. Data were normalized using a random forest algorithm-based signal correction method [20] available at (http://serrf.fiehnlab.ucdavus.edu).

### Assay 2 and 3. Hydrophilic interaction liquid chromatography (HILIC) Q-Exactive HF mass spectrometry for polar metabolites

#### Sample preparation

Metabolites were extracted from 20 **μ**L of mouse plasma using 1 mL of degassed, −20°C cold mixture of ACN:IPA:H2O(3:3:2, v/v/v). Samples were vortexed for 10 seconds, shaken for 5 minutes and then centrifuged for 2 minutes at 14,000 rcf. Two 450 **μ**L supernatant aliquots were transferred to new tubes. One tube was stored as a backup aliquot and another was dried in a SpeedVac concentrator. Sample were re-suspended with 100 **μ**L of ACN:H2O(80:20, v/v) which contained deuterium labeled internal standards (See Supplementary Table S18) prior to injection.

#### Data acquisition

3 μL sample aliquots were injected on a Waters Acquity UPLC BEH Amide column (150 mm length × 2.1 mm id; 1.7 μm particle size) maintained at 45°C. A Waters Acquity VanGuard BEH Amide pre-column (5 mm × 2.1 mm id; 1.7 μm particle size) was used as guard column. Mobile phase A was 100% LC-MS grade H2O with 10 mM ammonium formate and 0.125% formic acid and mobile phase B was 95:5 v/v ACN:H_2_Owith 10 mM ammonium formate and 0.125% formic acid. Gradient was started at 100% (B) for 2 min, 70% (B) at 7.7 min, 40% (B) at 9.5 min, 30% (B) at 10.25 min, 100% (B) at 12.75 min and isocratic until 16.75 min. The column flow was 0.4 mL/min. Vanquish UHPLC system (ThermoFisher Scientific) was used. A Thermo Q-Exactive HF Orbitrap MS instrument was operated in positive and negative ESI mdoes respectively with the following parameters: mass range 60-900 *m/z;* spray voltage 3.6kV (ESI+) and −3kV (ESI-), sheath gas (nitrogen) flow rate 60 units; auxiliary gas (nitrogen) flow rate 25 units, capillary temperature 320°C, full scan MS1 mass resolving power 60,000, data-dependent MSMS (dd-MSMS) 4 scans per cycle, normalized collision energy at 20%, 30% and 40%, dd-MSMS mass resolving power 15,000. Thermo Xcalibur 4.0.27.19 was used for data acquisition and analysis. Instruments was tuned and calibrated by manufacturer’s recommendations.

#### Data processing

Raw data files were converted to the mzML format using the ProteoWizard MSConvert utility. For each *m/z* values ion chromatogram was extracted with *m/z* thresholds of 0.005 Da and retention time threshold of 0.10 minute. Apex of the extracted ion chromatograph was used as peak height value and exported to a text file. Peak height files for all the samples were merged together to generate a data matrix. Targeted peak height signal extraction was performed using an R script which is provided at the GitHub repository (https://github.com/barupal). HILIC-POS data were not normalized because no batch effect was observed. HILIC-NEG data were normalized by the median value for each batch to remove batch effects.

### Assay 4 and 5. CSH-C18 Q-Exactive HF mass spectrometry for lipidomics

#### Sample preparation

Lipids were extracted from a 20 **μ**L of plasma. A 225 **μ**L of cold methanol (MeOH) containing a mixture of deuterated lipid internal standards (See Supplementary Table S18) was added and samples were vortexed for 10 seconds. Then 750 **μ**L of methyl tertiary-butyl ether (MTBE) was added. Samples were vortexed for 10 seconds and shaken for 5 mins at 4°C. Next, 188 **μ**L water was added and samples were vortexed for 10 seconds and centrifuged for 2 mins at 14000 rcf. Two 350 **μ**L aliquots from the non-polar layer were prepared. One aliquot was stored at −20 °C as a backup and the other was evaporated to dry in a SpeedVac. Dried extracts were resuspended using a mixture of methanol/toluene (9:1, v/v) (60 **μ**L) containing an internal standard [12-[[(cyclohexylamino)carbonyl]amino]-dodecanoic acid (CUDA)] used as a quality control. Method blanks and pooled human plasma (BioIVT) were prepared along with the study samples for monitoring the data quality.

#### Data acquisition

Extracted lipids were separated on an Acquity UPLC CSH C18 column (100 x 2.1 mm; 1.7 μm) maintained at 65°C. The mobile phases for positive mode consisted of 60:40 ACN:H_2_O with 10 mM ammonium formate and 0.1% formic acid (A) and 90:10 IPA:ACN with 10 mM ammonium formate and 0.1% formic acid (B). For negative mode, the mobile phase modifier was 10 mM ammonium acetate instead. The gradient was as follows: 0 min 85% (A); 0–2 min 70% (A); 2–2.5 min 52% (A); 2.5–11 min 18% (A); 11–11.5 min 1% (A); 11.5–12 min 1% (A); 12–12.1 min 85% (A); 12.1–15 min 85% (A). Sample temperature is maintained at 4°C in the autosampler. 2 **μ**L of sample was injected. Vanquish UHPLC system (ThermoFisher Scientific) was used. Thermo Q-Exactive HF Orbitrap MS instrument was operated in both positive and negative ESI modes respectively with the following parameters: mass range 120-1700 *m/z;* spray voltage 3.6kV (ESI+) and −3kV (ESI-), sheath gas (nitrogen) flow rate 60 units; auxiliary gas (nitrogen) flow rate 25 units, capillary temperature 320 °C, full scan MS1 mass resolving power 60,000, data-dependent MS/MS (dd-MS/MS) 4 scans per cycle, normalized collision energy at 20%, 30% and 40%, dd-MS/MS mass resolving power 15,000. Thermo Xcalibur 4.0.27.19 was used for data acquisition and analysis. The instrument was tuned and calibrated according to the manufacturer’s recommendations.

#### Data processing

Raw data files were converted to the mzML format using the ProteoWizard MSConvert utility. For each m/z values ion chromatogram was extracted with m/z thresholds of 0.005 Da and retention time threshold of 0.10 minute. Apex of the extracted ion chromatograph was used as peak height value and exported to a txt file. Peak height files for all the samples were merged together to generate a data matrix. Targeted peak height signal extraction was performed using an R script that is available at https://github.com/barupal. Extracted ion chromatograms for each peak were saved as pictures. CSH-POS and CSH-NEG data matrices were generated. No normalization was applied as minimum signal drift was observed during analysis.

### Assay 6 and 7. Bile acids-steroids, and oxylipin targeted analysis

#### Sample preparation

After thawing on ice and vortexing, 50 **μ**L of plasma from each sample was taken to polypropylene 96-well plate for extraction. The samples were spiked with internal standards of bile acids, steroids, and oxylipins at a concentration of 250 nM, resulting in a final concentration of 25 nM prior to LC-MS analysis. The suspensions were treated with antioxidant (0.2 ml/ml butylated hydroxytoluene and Ethylenediaminetetraacetic acid (EDTA)). 10 **μ**L of 1000 nM 1-cyclohexyluriedo-3-dodecanoic acid (CUDA) and 1-Phenyl 3-Hexadecanoic Acid Urea (PHAU) were added as quality markers for the analysis. ACN:MeOH 1:1 (v/v) were added to final volume of 250 μL. The samples were vortexed and incubated at 20°C for 30 min to precipitate protein. After centrifugation at 15,000 rcf for 5 min, the supernatant was transferred to a 0.2 μm PVDF filter plate (polyvinylidene fluoride membrane, Agilent). The solutions were collected in new polypropylene 96-well plates and stored in −20°C until analysis.

#### Data acquisition

For bile acids and steroids, reverse-phase liquid chromatography was performed on a Waters Acquity BEH C18 column (1.7 μm, 2.1×100 mm) with its corresponding Vanguard precolumn at 45 °C at a flow rate of 400 **μ**L/min. Mobile phase A was LC-MS grade H_2_O with 0.1% formic acid; mobile phase B was ACN with 0.1% formic acid. The 20 min gradient is: 0–0.5 min 10% B, 0.5–1 min 10–20% B, 1–1.5 min 20-22.5% B, 1.5–11 min 22.5–45% B, 11–12.5 min 45–95% B, 12.5–16 min 95% B, 16–16.5 min 95–10% B, 16.5–20 min 10% B.

For oxylipins, LC separation was conducted on the same column, but mobile phase A was H_2_O with 0.1% acetic acid and B was ACN:IPA 90:10 (v/v) with 0.1% acetic acid. The column was maintained at 45 °C at the flow rate of 250 **μ**L/min. A 16 min gradient was used with 0-1 min gradient from 25–40% B, 1–2.5 min 40–42% B, 2.5–4.5 min 42–50% B, 4.5–10.5 min 50–65% B, 10.5–12.5 min 65–75% B, 12.5–14 min 75–85% B, 14–14.5 min 85–95% B, 14.5–15 min 95–25% B, 15–16 min 25% B. Extracts were analyzed by liquid chromatography (Waters ACQUITY UPLC I-Class system) coupled to a Sciex 6500+ QTRAP hybrid, triple quadrupole linear ion trap mass spectrometer. 5 **μ**L of each extract was injected. Scheduled multiple reaction monitoring (MRM) was performed with optimized collision energies, de-clustering potentials, and collision cell exit potentials for individual analyte. A LC-MRM targeted method was used to analyze both bile acids and steroids with positive and negative polarity switching. Oxylipins were analyzed in another LC-MRM method in negative ionization mode only. All analytes were quantified against 6-point calibration curves using internal standards. Turbo Spray Ion Source parameters are: curtain gas (CUR) 25 psi, nebulizer gas (GS1) 50 psi, turbo-gas (GS2) 50 psi, electrospray voltage −4.5 kV/+3 kV, and source temperature 525 °C. Nitrogen was used as the collision gas. Software Analyst 1.6.3 and MultiQuant 3.0.2 (AB Sciex) were used for data acquisition and quantification. MRM transitions for the analytes are provided in the supplementary Table S8 and S9.

#### Data processing

MultiQuant version 3.0.2 was used for the peak integration and peak area computation. Peak integration settings were: Gaussian smooth width at 1.0 points, Retention half window at 10-15 sec, updated expected RT checkbox ‘NO’, minimum peak width at 8 points, minimum peak height at 750, noise at 40%, baseline subwindow at 1.7 minutes and peak splitting at 3 points. Multi-Quant software was also used for computing the molar concentrations for the analytes by using calibration curves created using internal standards as described in the supplementary file (Table S8 and S9).

### Data merging and filtering

Data matrices from each platform were combined to generate a joint dataset for all the samples. It contained a total of 1215 signals of identified metabolites (Table S12). Afterwards, signals were retained if relative standard deviation (RSD) was better than 50% and if fewer than 50% missing values were observed (Table S13). For metabolites that were detected in multiple platforms, data with the lowest relative standard deviation in the quality control samples were retained. The filtered dataset had 832 metabolites (Table S14). The simplified molecular-input line-entry system (SMILES) codes for all annotated lipids were obtained from the LipidBlast MSP file or from the PubChem Compound Identifier Exchange service (https://pubchem.ncbi.nlm.nih.gov/idexchange/idexchange.cgi) and provided in the data dictionary (Table S13). Chemical classes for the identified compounds were estimated using the ChemRICH software. Sample metadata is provided in the Table S15.

### Phenotype dataset

Mouse knockout’s phenotype data were downloaded from the IMPC database (www.mousephenotype.org) using their R-package IMPCData. First, allele accession numbers were matched to the IMPC database identifiers. Then, for each mouse accession, phenotype data were retrieved using the mouse strain identifier and phenotype identifiers (Table S1). The overall phenotype dataset is provided in the Table S2.

## 4. User Notes

Users can utilize raw spectra files, processed results, and the integrated metabolomics dataset. For the integration of phenotype and metabolomics dataset, the integrated dataset shall be used. Raw spectra files should be used to check the quality of detected peaks and to annotate unknown metabolites with new mass spectral libraries. Raw data files can be converted to mzML format for importing in other software such as mzR or MZ-Mine. Proper data transformation and scaling for each data matrix from the assays is recommended before performing univariate and multi-variate statistical analysis. The dataset is particularly interesting for researchers who focus on the biological functions of the 30 genes studied here, specifically, their potential roles in metabolism. The datasets given here present a new milestone in the field of metabolomics. We foresee its use in the developing next generation bioinformatics as well as in teaching courses for metabolomics and as test case for benchmarking software. As we have provided the annotation database, mass spectral libraries, and protocol details, these resources can be used to re-create similar datasets for other cohorts of the blood specimens. We performed a ChemRICH class annotation for the structurally identified compounds and found that almost 80 chemical classes are covered. These chemical groups can be associated with genes and with phenotypes.

## Supplementary Materials

The following are available online at www.mdpi.com/link,

Table S1 Phenotype details

Table S2 Phenotype data for knockouts

Table S3 WCMC – GCMS annotation database

Table S4 WCMC – HILIC annotation database (ESI POS)

Table S5 WCMC – HILIC annotation database (ESI NEG)

Table S6 WCMC – CSH annotation database (ESI POS)

Table S7 WCMC – CSH annotation database (ESI NEG)

Table S8 WCMC – Targeted metabolomics database (Bile Acids and Steroids)

Table S9 WCMC – Targeted metabolomics database (Oxylipins)

Table S10 Sample and file name mapping

Table S11 Raw metabolomics dataset for mouse knockouts

Table S12 Filtered metabolomics dataset

Table S13 Data Dictionary

Table S14 Data Matrix

Table S15 Sample metadata

Table S16 Reagents and material used

Table S17 Analysis sequences for all assays

Table S18 Internal standards for LCMS assays

## Acknowledgments

The study was funded by the “West Coast Metabolomics Center for Compound Identification” was provided by the National Institutes of Health under the award number NIH U2C ES030158 (to O.F.). We thank staff members of the West Coast Metabolomics Center, the International Mouse Phenotyping Center and the Mouse Metabolic Phenotyping Center for their support in implementing the project.

## Author Contributions

All authors contributed in writing the manuscript. O.F. and D.B. conceptualized the study. D.B generated the consolidated metabolomics dataset. D.B. and S.F. performed statistical analysis. Y.Z and Dinesh Barupal retrieved the phenotype data. T.S., G.B. and P.F. and Y.C. did targeted Assays for Bile acids, Steroids and Oxylipins. B.S.R., B.H.,T.S., C.S.B., J.S.F, M.R.S and B.W did LCMS data acquisition. C.S.B, J.S.F and D.B. processed LCMS data. L.V. and O.F. processed GC-TOFMS data. T.K., M.R.S. and A.V. contributed LC-MS/MS mass spectral libraries

